# MFGE8 inhibits insulin signaling through PTP1B

**DOI:** 10.1101/2023.05.30.542928

**Authors:** Ritwik Datta, Michael J Podolsky, Christopher D Yang, Diana L. Alba, Sukhmani Singh, Suneil Koliwad, Carlos O Lizama, Kamran Atabai

**Author notes:** **Corresponding author:** Kamran Atabai, MD.

## Abstract

The role of integrins in regulating insulin signaling is incompletely understood. We have previously shown that binding of the integrin ligand milk fat globule epidermal growth factor like 8 (MFGE8) to the αvβ5 integrin promotes termination of insulin receptor signaling in mice. Upon ligation of MFGE8, β5 complexes with the insulin receptor beta (IRβ) in skeletal muscle resulting in dephosphorylation of IRβ and reduction of insulin-stimulated glucose uptake. Here we investigate the mechanism by which the interaction between β5 and IRβ impacts IRβ phosphorylation status. We show that β5 blockade inhibits and MFGE8 promotes PTP1B binding to and dephosphorylation of IRβ resulting in reduced or increased insulin-stimulated myotube glucose uptake respectively. The β5-PTP1B complex is recruited by MFGE8 to IRβ leading to termination of canonical insulin signaling. β5 blockade enhances insulin-stimulated glucose uptake in wild type but not *Ptp1b* KO mice indicating that PTP1B functions downstream of MFGE8 in modulating insulin receptor signaling. Furthermore, in a human cohort, we report serum MFGE8 levels correlate with indices of insulin resistance. These data provide mechanistic insights into the role of MFGE8 and β5 in regulating insulin signaling.

## INTRODUCTION

Milk fat globule epidermal growth factor like 8 (MFGE8) is a secreted integrin ligand that regulates whole body metabolism through effects on nutrient absorption (1,2), gastrointestinal motility (1), and lipid homeostasis (3,4). In mice, MFGE8 impacts glucose homeostasis in multiple ways. Global *Mfge8* KO mice develop insulin resistance concomitant with the development of differences in body fat composition at 10 weeks of age (2). In contrast, we have recently reported that MFGE8 can negatively regulate insulin receptor signaling acutely (5). Antibody-mediated blockade of MFGE8 or the αvβ5 integrin, the cell surface receptor for MFGE8, leads to enhanced insulin sensitivity in wild type (WT) mice (5). However, a detailed molecular understanding of the effect of MFGE8 and intermediaries on the insulin receptor is still lacking.

In humans, a growing body of literature links MFGE8 with insulin resistance. Serum MFGE8 levels are increased in patients with type II diabetes mellitus (T2D) and correlated with the degree of hemoglobin glycosylation (6,7). Serum MFGE8 levels correlate with markers of insulin resistance in two diabetic cohorts of patients from China, one with gestational diabetes and one with T2D mellitus (8,9). More recently, a missense mutation found in Punjabi Sikhs increases circulating MFGE8 and markedly increases the risk of developing T2D in this population (10,11). These latter data are strongly suggestive of a causal relationship in humans between MFGE8 and glucose homeostasis.

Insulin signaling is tightly regulated through phosphorylation of intermediaries in the insulin receptor pathway. Insulin binding to the α subunit of the insulin receptor (IRα) induces a conformational change that activates intrinsic receptor tyrosine kinase activity resulting in auto-phosphorylation and activation of the insulin receptor β subunit (IRβ) (12,13). Tyrosine kinase activity of activated IRβ in turn triggers a cascade of phosphorylation events in downstream mediators ultimately inducing increased translocation of glucose transporter type 4 (GLUT4) to the plasma membrane and increased glucose uptake (14,15). The activity of protein tyrosine phosphatases opposes that of protein tyrosine kinases thereby dampening insulin receptor signaling (16).

As referenced above, we recently described an autoregulatory pathway through which insulin increases outer cell membrane MFGE8 levels. MFGE8 then binds the αvβ5 integrin leading to increased co-localization and binding of β5 to IRβ. This interaction promotes dephosphorylation of IRβ and reduces skeletal muscle glucose uptake in response to insulin (5). The effect of MFGE8-β5 on insulin receptor phosphorylation could be the result of reduced kinase or increased phosphatase activity. In our previous work, antibody-mediated blockade of β5 leads to delayed recovery of serum glucose levels after insulin challenge in mice (5), similar to what has been reported with genetic deletion of the phosphatase Protein-tyrosine phosphatase 1B (*Ptp1b*) in mice (17). PTP1B binds to and dephosphorylates IRβ and insulin receptor substrate 1 (IRS1) promoting termination of the insulin signaling cascade (18-21). Dysregulation of PTP1B activity has been implicated in insulin resistance and T2D (18,22) and inhibition of PTP1B improves insulin sensitivity, glucose uptake, and glucose homeostasis in preclinical models (18,22). In the present work we investigate the mechanism of MFGE8-β5-mediated inhibition of insulin signaling, and whether PTP1B functions downstream of MFGE8 and αvβ5 in this process. We show here that β5 is found in complex with PTP1B and that MFGE8 binding to β5 induces recruitment of this complex to IRβ thereby causing IRβ dephosphorylation and deactivation of the canonical pathway for insulin-mediated glucose uptake.

## RESULTS

### β5 blockade prevents the effect of insulin on PTP1B phosphatase activity

In our recently published work (5), we showed enhanced in vivo skeletal muscle insulin receptor phosphorylation after insulin treatment in the presence of a β5 blocking antibody. However, the mechanism of this enhanced phosphorylation was not delineated. Therefore, we first evaluated whether β5 blockade impacts PTP1B phosphatase activity in mice. We immunoprecipitated PTP1B from skeletal muscle lysates treated with insulin in the presence of a β5 blocking or isotype control antibody and measured phosphatase activity of the immuno-precipitate on a tyrosine phospho-peptide substrate. In control samples insulin treatment reduced PTP1B activity 15 minutes after administration which then recovered to baseline levels at 60 minutes consistent with what has been reported in the literature (23). In the presence of β5 blockade, insulin induced a similar drop in PTP1B activity 15 minutes after administration. However, there was no recovery of PTP1B activity at the 60-minute time point with β5 blockade (Figure 1A).

**Figure 1:**
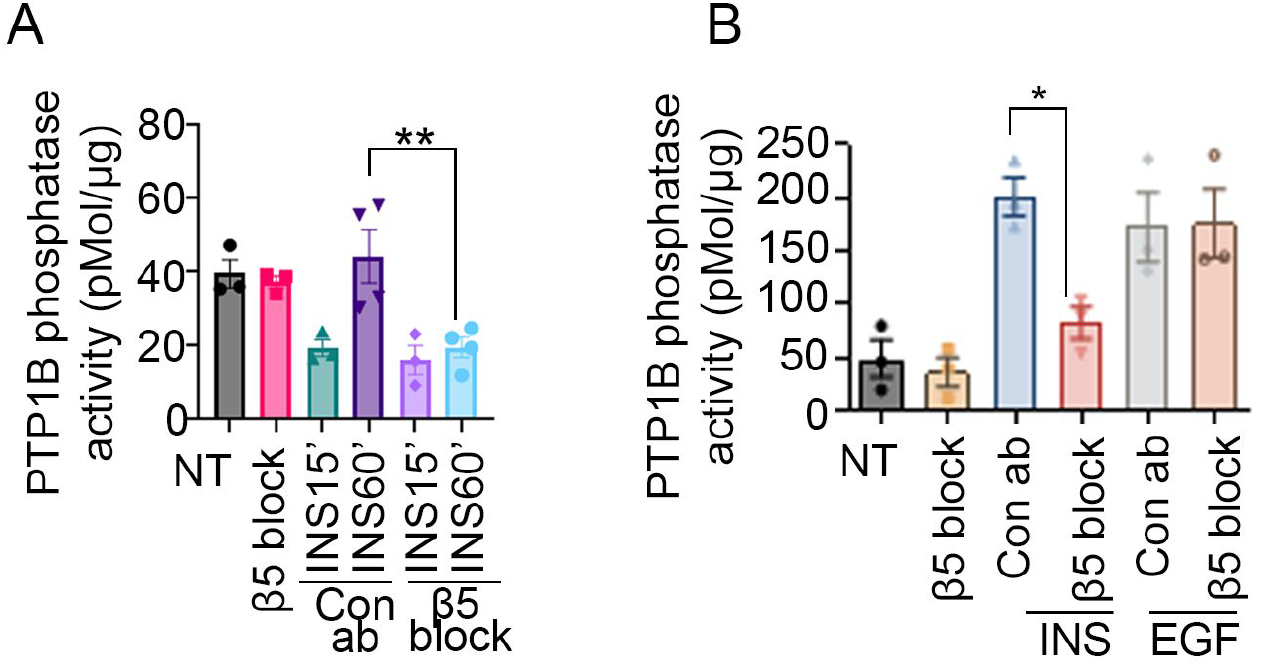
β5 blockade reduces insulin-stimulated PTP1B phosphatase activity. (A,B) PTP1B phosphatase activity in the presence of β5 blocking or isotype control antibody in (A) skeletal muscle lysates harvested from WT mice 15 and 60 minutes after IP insulin (1U/kg) administration and (B) in Hela cells 30 minutes after EGF or insulin (100 nm) treatment. N=4 male mice in each group for panel A. Data merged from 2 independent experiments. N=3 independent experiments for panel B. Data are expressed as mean ± SEM; ∗p < 0.05, ∗∗p < 0.01, and analyzed by One-way ANOVA followed by Bonferroni’s post-test.

We next evaluated whether altered PTP1B activity with β5 blockade is specific for insulin-mediated signaling. In addition to the insulin receptor, PTP1B dephosphorylates tyrosine residues of the Epidermal Growth Factor Receptor (EGFR) (24,25). Therefore we used Hela cells, which express both the EGFR (26) and the insulin receptor (27), to measure PTP1B phosphatase activity in the presence of β5 blocking or isotype control antibody after treatment with insulin or EGF. β5 blockade inhibited PTP1B activity stimulated by insulin but not EGF (Figure 1B) indicating that β5 specifically modulates PTP1B phosphatase activity associated with insulin signaling.

### The β5 integrin is in complex with PTP1B

We next evaluated whether β5 integrin forms a complex with PTP1B via co-immunoprecipitation. We administered β5 blocking or isotype control antibody to WT mice 1 hour before IP insulin treatment and then performed co-immunoprecipitation studies to measure β5-PTP1B binding from hind-leg skeletal muscle lysates of WT mice 1 hour after insulin treatment. Lysates were immunoprecipitated with an antibody targeting PTP1B followed by Western blot for β5 or IRβ or with an antibody targeting β5 followed by Western blot for PTP1B or IRβ. Both approaches showed that β5 and PTP1B forms a complex at baseline. Insulin treatment increased co-immunoprecipitation of β5 and PTP1B which was further increased after pre-treatment with the β5 blocking antibody (Figure 2A). While the association between PTP1B and IRβ was increased, as expected, by insulin (28), this effect was abrogated in the presence of β5 blockade (Figure 2A). These data are consistent with our previously published work showing reduced skeletal muscle insulin-stimulated interaction between β5 and IRβ in the presence of β5 blockade (5).

**Figure 2:**
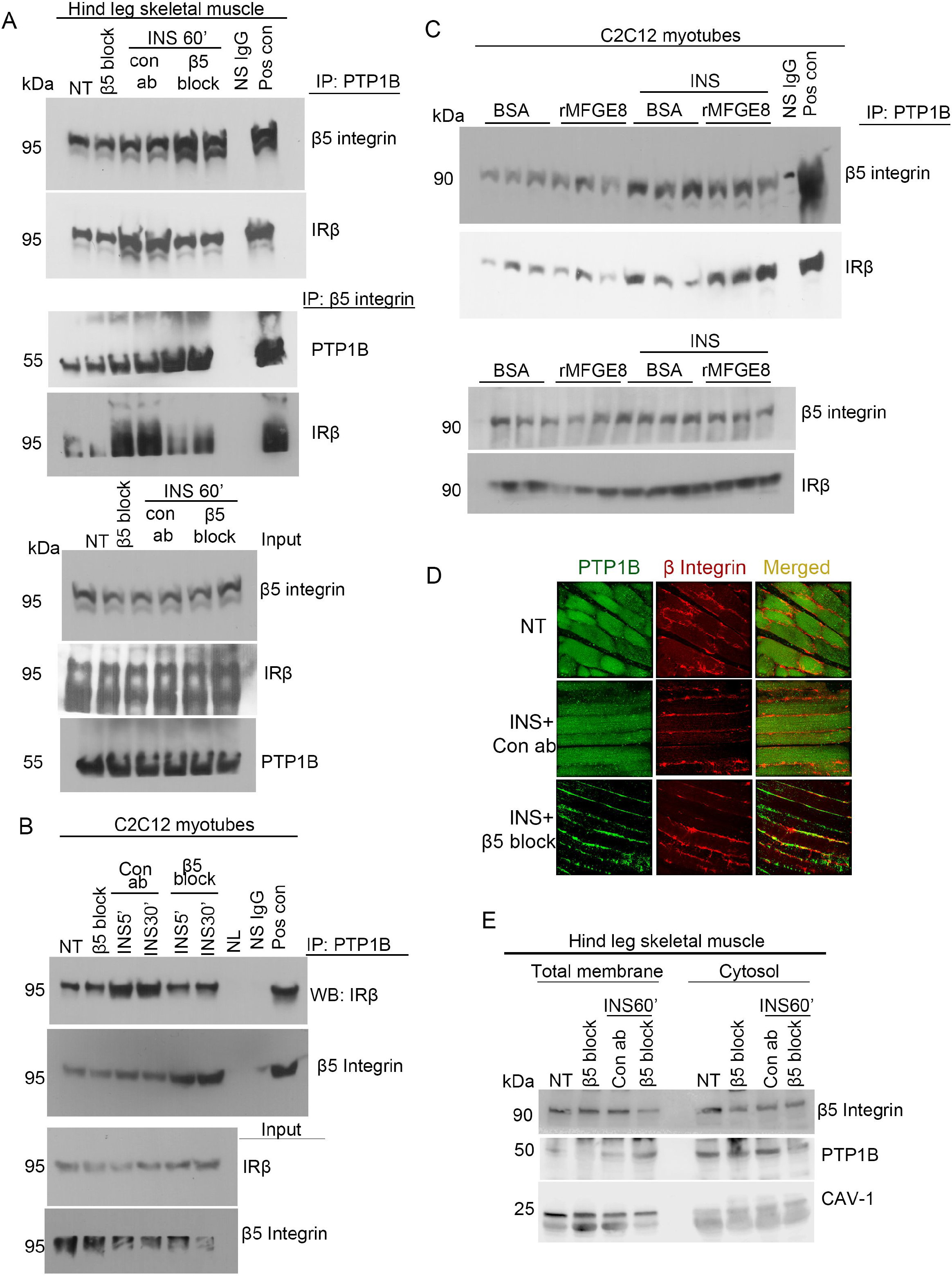
β5 complexes with PTP1B and IRβ. (A) Co-immunoprecipitation studies showing interaction of PTP1B with β5 and IRβ in hind leg skeletal muscle lysates from mice in presence of β5 blocking or isotype control antibody 60 minutes after IP insulin (1U/Kg) treatment. Blots represent 2 independent experiments with N=4 male mice for insulin-treated groups and 2 mice without insulin treatment. (B,C) Co-immunoprecipitation studies showing interaction of PTP1B with β5 and IRβ in C2C12 myotubes treated with (B) β5 blocking (5μg/mL) or isotype control antibody or (C) MFGE8 (10 μg/mL) or BSA control in the presence and absence of insulin for 30 minutes. N=4 male mice in total. Panel B represents 2 independent experiments. For panel C, cell culture, treatment and protein isolation performed on 3 different days. After immunoprecipitation, protein samples were run on the same gel for western blot. (D) Immunostaining of PTP1B (green) and β5 integrin (red) in skeletal muscle cross sections from mice treated with insulin for 60 minutes in presence of β5 blocking or isotype control antibody. Images are representative from 2 independent experiments. (E) Cell fractionation of insulin-treated skeletal muscle tissue samples from mice pretreated with β5 blocking or isotype control antibody and insulin for 60 minutes, followed by Western blot for PTP1B and β5 in the cytoplasmic and membrane fraction. Western blotting for CAVEOLIN-1 (CAV-1) confirmed enrichment of the membrane fractions. N=1 per group per experiment. Western blots are representative of 3 independent experiments.

We validated these findings in vitro by treating C2C12 myotubes with β5 blocking or isotype control antibody in the presence of insulin for 5 and 30 minutes, immunoprecipitating lysates with an antibody against PTP1B, and subsequently performing Western blot for β5. Consistent with our in vivo findings (Figure 2A), β5 blockade enhanced the interaction between PTP1B and β5 while reducing the interaction between PTP1B and IRβ in the presence of insulin at both time points (Figure 2B). These data show that PTP1B complexes with β5 integrin, consistent with the functional interaction suggested by Figure 1.

We have previously shown that the effect of β5 integrin on insulin signaling is specific for MFGE8 and not induced by other β5 ligands such as fibronectin or vitronectin (5). To determine whether the β5-PTP1B interaction is modified by MFGE8, we treated C2C12 myotubes with recombinant MFGE8 (rMFGE8) with and without insulin for 30 minutes and performed co-immunoprecipitation of cell lysates with an antibody targeting PTP1B followed by western blot for β5 and IRβ. rMFGE8 treatment did not impact the association between β5 and PTP1B at baseline or after insulin treatment (Figure 2C). However, in the presence of insulin rMFGE8 increased the interaction between IRβ and PTP1B (Figure 2C). Taken together, these data indicate that MFGE8 increases recruitment of the β5-PTP1B complex to IRβ.

We next performed immunofluorescence staining for β5 and PTP1B in sections from hind-leg skeletal muscle of mice treated with insulin for 60 minutes in the presence of either β5 blocking or isotype control antibody. We found that β5 and PTP1B co-localized (Figure 2D) and that co-localization increased after insulin treatment in the presence of β5 blocking antibody consistent with our co-IP experiments (Figure 2D). Interestingly, there was a striking enrichment of membrane localization of PTP1B after insulin treatment in the presence of β5 blockade (Figure 2D).

To further investigate the subcellular localization of PTP1B and β5 integrin, we performed cell fractionation of skeletal muscle lysates harvested from the hind-legs of WT mice 60 minutes after insulin treatment and pretreated with β5 blocking or isotype control antibody. Consistent with our immunofluorescence data, β5 blockade led to increased membrane and reduced cytosolic PTP1B expression after insulin treatment (Figure 2E). Taken together, these data suggest that by preventing MFGE8 ligation of β5, β5 blockade leads to sequestration of PTP1B in complex with the integrin at the cell membrane in a manner that reduces recruitment of the complex to IRβ, which functionally decreases PTP1B-mediated dephosphorylation of IRβ.

### β5 blockade promotes insulin-stimulated AKT signaling

To determine how MFGE8 and β5 impact canonical insulin signaling, we pretreated C2C12 myotubes with β5 blocking antibody for 1 hour, treated with insulin for 5 or 30 minutes, and evaluated AKT phosphorylation at serine 473 (ser473) residue by western blot. β5 blockade augmented insulin-stimulated AKT phosphorylation at both timepoints (Figure 3A-B). Additionally, treatment with rMFGE8 dampened insulin-mediated AKT phosphorylation (Figure 3C-D). To validate these findings in vivo, we administered β5 blocking or isotype control antibody to WT mice 1 hour before treating them with insulin (5, 15, or 30 minutes) and assessed ser473 AKT phosphorylation in hind-leg skeletal muscle protein lysates. β5 blockade enhanced AKT phosphorylation at the 5- and 30-minute timepoints as compared with control antibody-treated mice (Figure 3E-F). We interpret these data to indicate that β5 blockade causes enhanced and persistent activation of canonical insulin signaling pathway.

**Figure 3:**
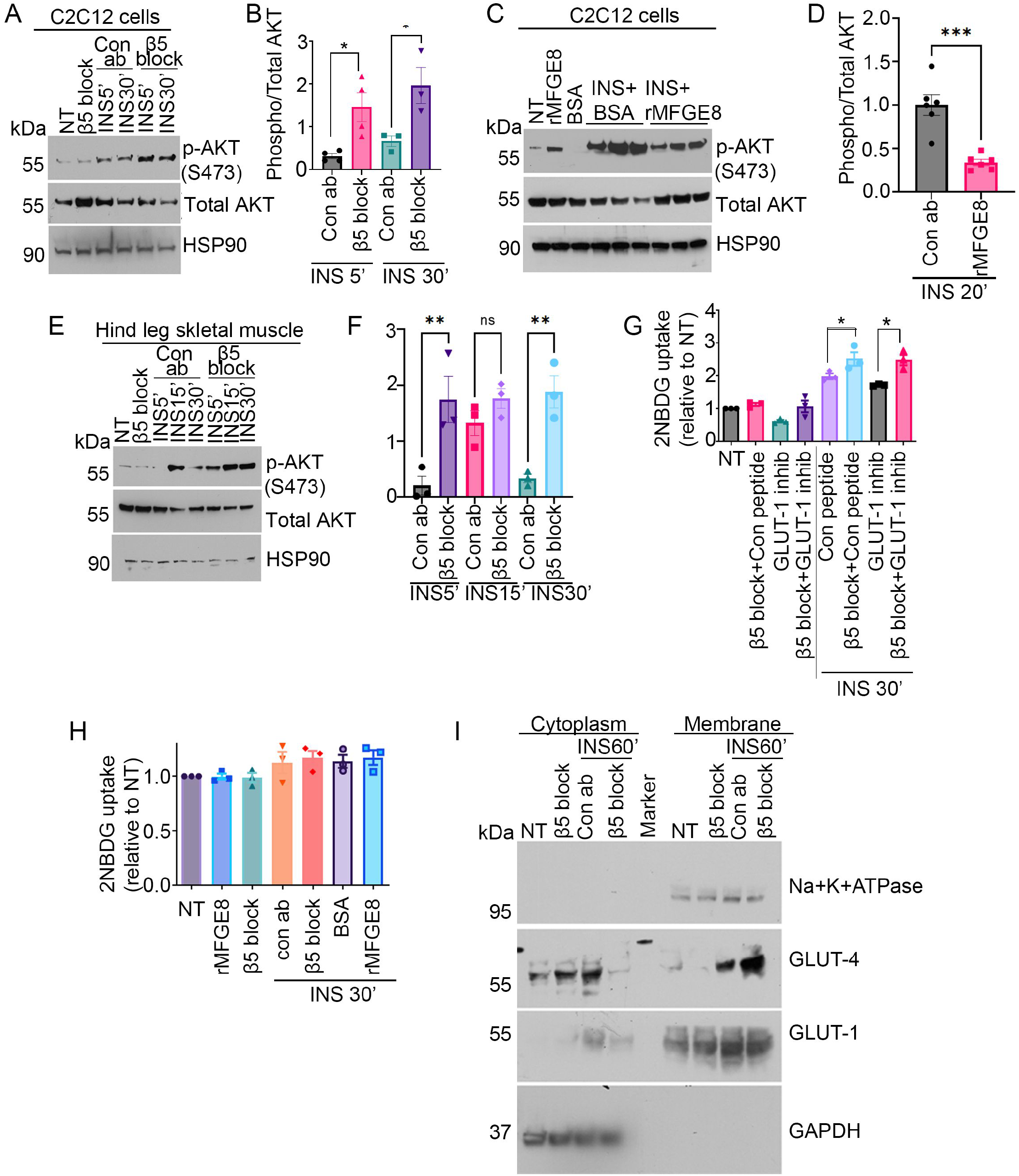
β5 blockade activates canonical insulin signaling. (A) Representative western blot showing phosphorylated (serine 473 residue, S473) and total AKT levels in C2C12 myotubes in the presence of β5 blocking or isotype control antibody treated with insulin for 30 minutes. N=3 independent experiments. (B) Densitometric analysis of western blots from panel A. (C-D) Representative western blot showing phosphorylated AKT and total AKT levels in C2C12 myotubes treated with rMFGE8 (10μg/mL) and insulin for 20 minutes. N=6 independent samples from 2 independent experiments. (D) Densitometric analysis of western blots from panel C. (E) Representative western blot of phosphorylated and total AKT levels in skeletal muscle lysates from mice treated with insulin for 5, 15 and 30 minutes in presence of β5 blocking or isotype control antibody. N=1 male mouse per group per experiment with 3 independent experiment. (F) Densitometric analysis of western blots from panel E. (G) Effect of GLUT-1 inhibitor (1μM) or control peptide on 2NBDG uptake in C2C12 myotubes treated with β5 blocking or isotype control antibody after insulin stimulation. N=3 independent experiments. Data expressed as relative fold change to untreated cells (NT). (H) Effect of β5 blockade (5μg/mL) or rMFGE8 treatment (10 μg/mL) on 2NBDG uptake in 3T3 fibroblasts in the presence or absence of insulin. N=3 independent experiments. (I) Western blot showing GLUT-1 and GLUT-4 levels in cytosolic and membrane fractions isolated from C2C12 myotubes treated with insulin for 30 minutes in presence of β5 blocking or isotype control antibody. Na^+^K^+^ATPase and GAPDH were used as loading controls for membrane and cytosolic fractions respectively. The western blot represents 3 independent experiments. Data are expressed as mean ± SEM; ∗p < 0.05, ∗∗p < 0.01, ∗∗∗p < 0.001, and analyzed by One-way ANOVA followed by Bonferroni’s post-test.

### β5 blockade increases glucose uptake through glucose transporter 4

Next, we evaluated how β5 blockade impacts glucose transporter activity. Skeletal muscle predominantly utilize glucose transporter 1 (GLUT-1) and glucose transporter 4 (GLUT-4) for glucose uptake (14,29,30). We treated C2C12 myotubes with a small molecule inhibitor of GLUT-1 in the presence of β5 blocking or isotype control antibody. The ability of β5 blockade to augment insulin-mediated glucose uptake in these cells was unaffected by GLUT-1 inhibition (Figure 3G). Therefore we then evaluated whether β5 blockade impacts insulin-stimulated glucose uptake in 3T3 fibroblasts, a cell line that expresses GLUT-1 (31) but not GLUT-4 (32). β5 blockade had no effect on insulin-stimulated glucose uptake in these cells suggesting that the effect of β5 is independent of GLUT-1 (Figure 3H). We next performed cell fractionation and assessed cytoplasmic and membrane levels of GLUT-1 and GLUT-4 in insulin-treated C2C12 myotubes in the presence of β5 blocking or isotype control antibody. β5 blockade markedly increased membrane enrichment of GLUT-4 without an appreciable effect of membrane GLUT-1 expression in the presence of insulin (Figure 3I). Taken together, these data indicate that β5 blockade impacts insulin-stimulated glucose uptake in myotubes by promoting GLUT-4 movement to the cell membrane.

### β5 dampens insulin signaling via PTP1B

To determine whether PTP1B is necessary for the effect of β5 integrin on insulin signaling, we measured insulin-mediated glucose uptake in *Ptp1b* KO myotubes after treating cells with β5 blocking or isotope control antibody. *Ptp1b* KO myotubes showed enhanced insulin-mediated glucose uptake compared to WT myotubes that was not affected by β5 blockade (Figure 4A). To validate our findings in vivo, we administered β5 blocking or isotype control antibody to WT and *Ptp1b* KO mice and performed a glucose tolerance test. *Ptp1b* KO mice had significantly reduced blood glucose levels after IP glucose challenge as compared with WT mice. Antibody mediated blockade of β5 in *Ptp1b* KO mice did not further impact blood glucose level (figure 4B). Taken together these data show no additive effect on insulin-stimulated skeletal muscle glucose uptake with simultaneous silencing of *Ptp1b* and β5 and support the hypothesis that PTP1B functions downstream of MFGE8 and β5 in modulating insulin signaling (Figure 5) (5).

**Figure 4:**
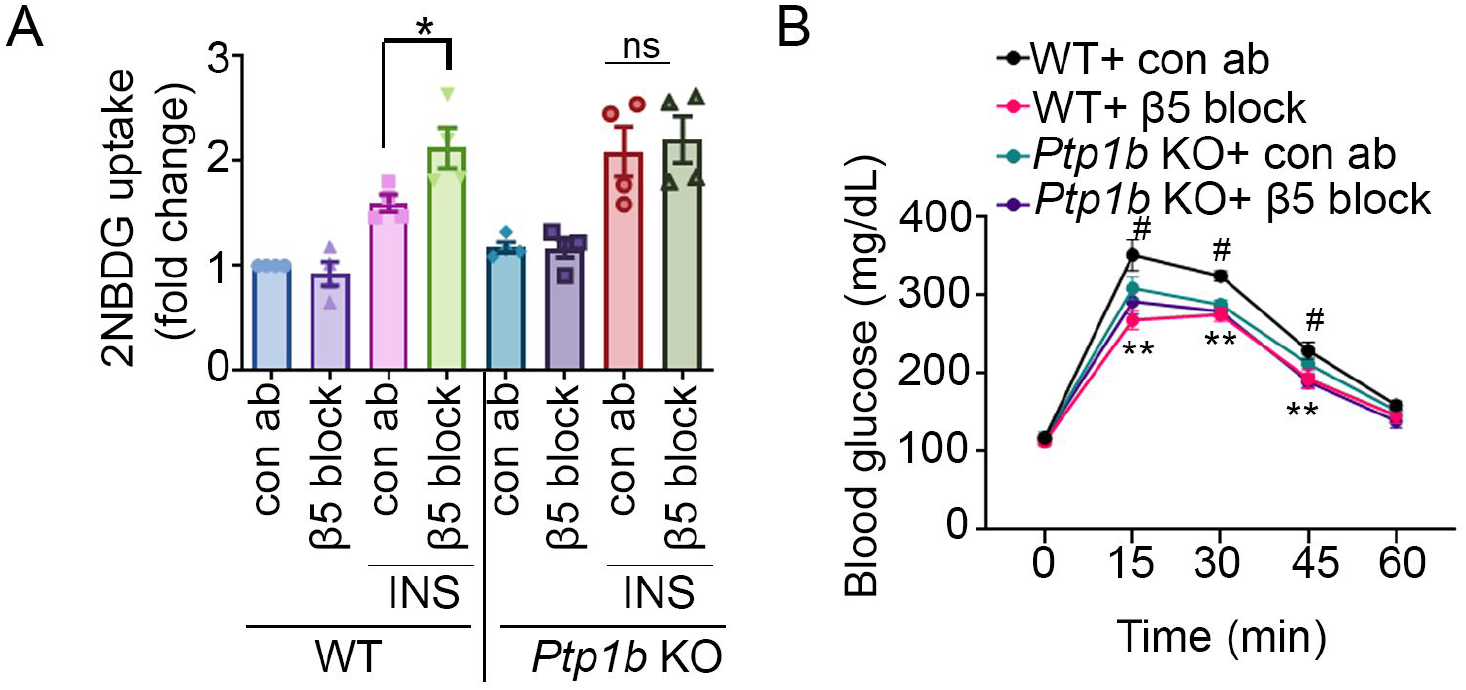
β5 blockade impacts insulin signaling through PTP1B. (A) 2NBDG uptake assay in WT and *Ptp1b* KO myotubes in presence of β5 blocking or isotype control antibody and insulin. N=4 independent experiments. (B) Glucose tolerance test in 7 to 8-week-old male WT and *Ptp1b* KO mice after IP injection of β5 blocking or isotype control antibody. N=6 male mice per group from 2 independent experiments are presented. Data are expressed as mean ± SEM. Data in panel (A) were analyzed by One-way ANOVA followed by Bonferroni’s post-test. ∗p < 0.05. Data in panel (B) were analyzed by 2-way ANOVA followed by Tukey’s post-test. **p<0.01, *p < 0.05, when comparing WT+Con ab versus WT+β5 blockade groups; #p<0.05 when comparing WT+Con ab versus *Ptp1b*KO+Con ab groups.

**Figure 5:**
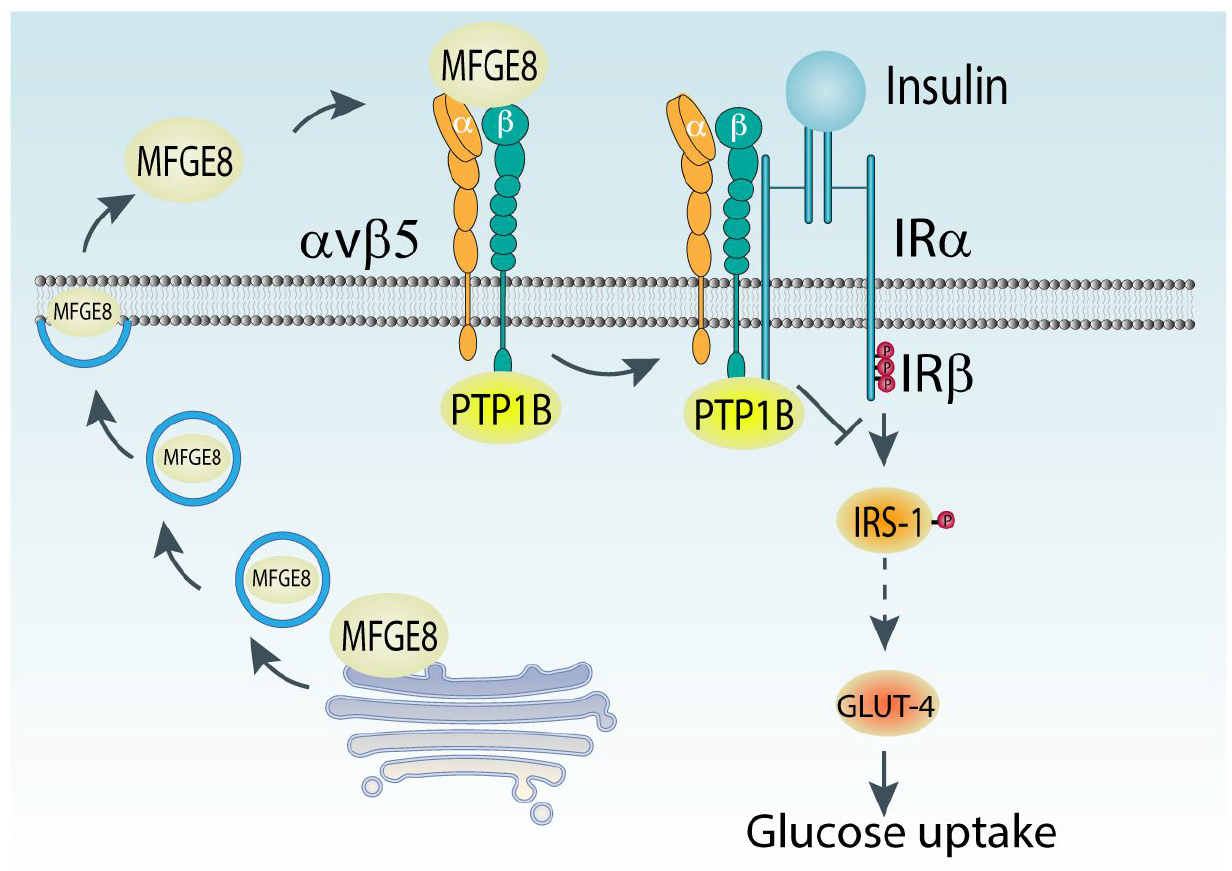
Model depicting how MFGE8 impacts insulin signaling through PTP1B. MFGE8 binding of the β5 integrin on the outer cell membrane leads to recruitment of the β5-PTP1B complexes to IRβ. PTP1B subsequently dephosphorylates IRβ leading to reduced translocation of GLUT4 to the cell surface and reduced glucose uptake in response to insulin.

### Physiological regulation of PTP1B/β5 interaction

We have previously shown that fasting decreases while refeeding increases serum MFGE8 and insulin levels and the association of IRβ and β5 in skeletal muscle (5). To examine whether refeeding increases the association of IRβ and PTP1B, we immunoprecipitated PTP1B from skeletal muscle lysates from fasted and refed mice and performed western blot for IRβ and β5. Refeeding increased the association between PTP1B and IRβ as well as interactions between PTP1B and β5 as compared with the fasted state (Figure 6) indicating that these interactions are regulated by nutritional status, further highlighting the role of this pathway during physiological conditions.

**Figure 6:**
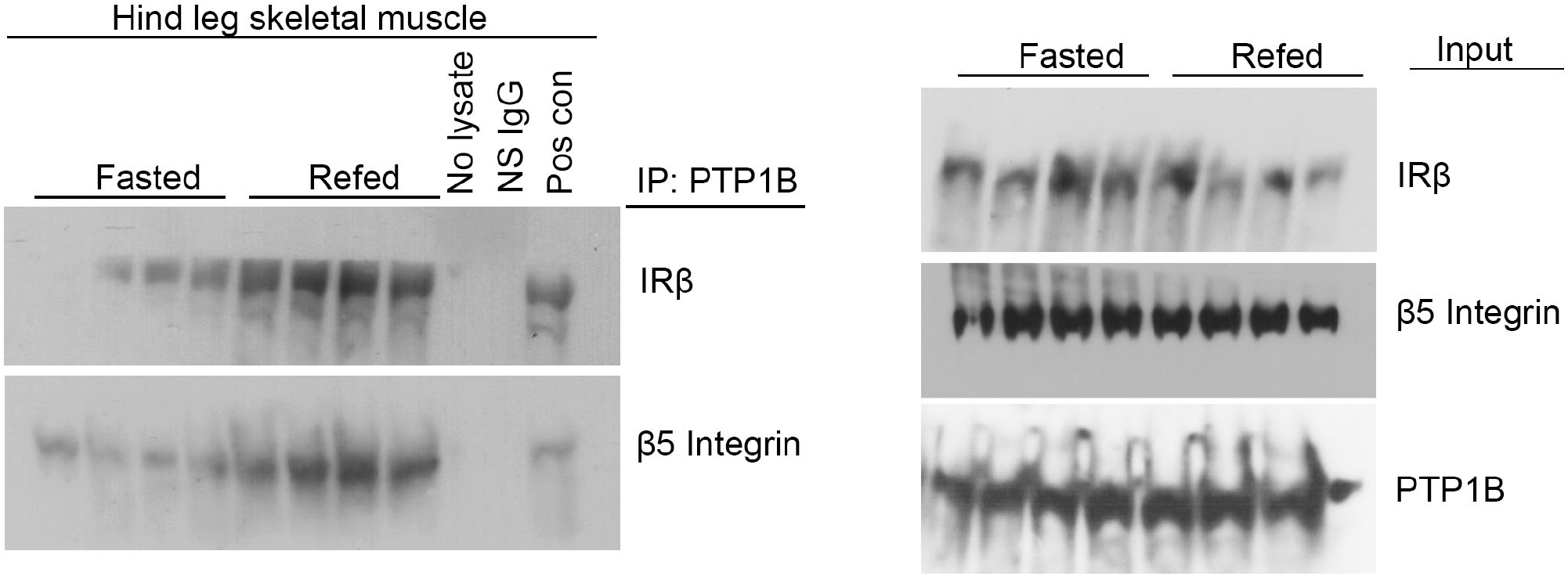
Physiological regulation of PTP1B/β5 interaction. Co-immunoprecipitation studies showing interaction of PTP1B with β5 and IRβ in hind-leg skeletal muscle lysates from mice fasted for 16h or mice refed for 1h after a 16h fast. N=4 mice in each group.

### Correlation between serum Mfge8 and insulin resistance in human subjects

Finally we performed analyses in human data, providing additional relevance to our findings. As above, serum MFGE8 levels are correlated with indices of insulin resistance in Chinese cohorts of patients with T2D (8) or gestational diabetes (9). To expand on these findings, we examined serum MFGE8 levels and indices of insulin resistance in a multi-ethnic cohort of 25-to 65-year-old healthy and primarily White, Chinese, and Hispanic individuals living in the San Francisco Bay Area (Supplementary Table 1). We performed multivariate linear regression analysis utilizing significant independent predictor variables determined by Spearman correlation coefficients (Supplementary Table 2) to identify independent predictors of insulin resistance as measured by the Homeostatic Model Assessment of Insulin Resistance (HOMA-IR). MFGE8 was a significant independent predictor of insulin resistance in this cohort on par with what we found for blood glucose, fasting glucose, and insulin levels (Supplementary Table 3).

## Discussion

Despite many decades of work, there is still much unknown about insulin signaling and glucose homeostasis. The regulation of metabolic homeostasis is enormously complex, but elucidating this complexity can yield insights not just about normal physiology, but also about disease states. The ongoing global pandemic of obesity and one of its key complications, T2D, highlight that there is still urgent unmet medical need in understanding all pathways of regulation of glucose and insulin signaling.

We recently identified an autoregulatory pathway by which insulin promotes termination of insulin receptor signaling by increasing the amount of MFGE8 at the outer cell membrane (5). MFGE8 binds the αvβ5 integrin, increasing the association of αvβ5 with the insulin receptor and resulting in insulin resistance through reduced phosphorylation/activation of the insulin receptor signaling pathway (5). In the current work, we build on these findings and show that decreased insulin receptor phosphorylation induced by MFGE8 and αvβ5 occurs via the phosphatase PTP1B. Our data supports a model wherein β5, PTP1B, and IRβ are in a complex that is modulated by MFGE8 and β5. MFGE8 interaction with β5 increases the association between PTP1B and IRβ. By contrast, our co-IP studies indicate that blockade of the integrin in the presence of insulin increases the association of β5 with PTP1B while decreasing the association with IRβ. Our immunofluorescence and cell fractionation studies validate these findings by showing increased membrane localization of PTP1B in the presence of insulin and integrin blockade suggesting that ligand (MFGE8) binding to β5 releases the β5-PTP1B complex from the membrane such that it can strengthen its association with IRβ. PTP1B is known to be important in insulin receptor desensitization (19) – increased PTP1B expression is associated with insulin resistance (17,33) and PTP1B is being evaluated as a therapeutic for patients with T2D (34). Our data add to the field by demonstrating a mechanism that regulates PTP1B activity during insulin signaling. Furthermore, our data suggest that biological manipulation of β5 may provide an alternate or additional mechanism to inhibit PTP1B that could be of therapeutic value.

One limitation of our work is that we cannot rule out the potential impact of β5 blockade on the intrinsic tyrosine kinase activity of the insulin receptor independent of its effect on PTP1B. Additionally, we have not explored whether increasing the association of β5 with IRβ impacts insulin receptor structure such that it is less receptive to insulin binding thereby reducing the strength of insulin signaling.

There is an increasing body of literature in human subjects supporting the role of MFGE8 in diabetes mellitus. From our perspective, the most intriguing data comes from work showing that a missense mutation in MFGE8 dramatically increases the risk for developing T2D in Punjabi Sikhs, a population prone to developing non-obese T2D (10). Additional studies of interest include two cohorts of patients from China, one with T2D and one with gestational diabetes, in whom serum MFGE8 levels correlate with multiple indices of insulin resistance (8,9). Interestingly, in the T2D cohort, the association is prominent in non-obese subjects and absent in obese subjects (8). We were therefore curious to examine this relationship in a North American cohort consisting of similar proportion of healthy White, Chinese, and Hispanic individuals from the general population. Our findings are consistent with the previously published correlation of serum MFGE8 levels with indices of insulin resistance (8,9). In our cohort, serum MFGE8 levels significantly predict HOMA-IR on par with what we observed for fasting glucose and insulin levels. Interestingly, this correlation was not restricted to non-obese patients in contrast to what is been reported in the Chinese cohort (8). Despite relatively modest sample sizes, these 4 cohorts (8-10) validate our mechanistic work in mice indicating a role for MFGE8 in inducing insulin resistance (2,5) and lay the foundation for therapeutic targeting of this pathway to ameliorate insulin resistance.

A better understanding of pathobiology and etiology of insulin resistance can open the door to new therapeutic avenues to treat patients. Although insulin therapy is the mainstay of treatment for type 1 diabetes mellitus and is a major component of therapeutic regimes for patients with T2D, chronic insulin treatment can have numerous deleterious effects. For example, insulin can promote weight gain, a particularly disadvantageous effect for obese patients with T2D. Insulin therapy in insulin resistant patients can require the administration of very high doses of exogenous insulin raising the risk of serious consequences from dosing errors. Therefore approaches that leverage endogenous pathways to enhance insulin signaling could be one means to avoid some of these problems. Therefore, we believe our work is an important addition to understanding foundational mechanisms of insulin signaling regulation.

## Materials and methods

### Mice

All studies utilized 6-10 week-old male age-matched mice in C57/bl6 background. *Ptp1b* KO mice are in the C57/bl6 background and have been extensively characterized (17,35).

### Cell culture and treatments

C2C12 myoblasts (ATCC) were cultured in DMEM supplemented with 10% FBS at 37°C and 5% CO_2_. Cells were then passaged and differentiated following previously published protocol (5). The experiments were performed after 4 days of differentiation. C2C12 cells were serum-starved for 4 h. Treatment with β5 blocking (5 μg/mL) antibody or a control antibody was administered 1 h before insulin (100 nm) stimulation. rMFGE8 (10 μg/mL) or BSA control was added to the media concurrently with insulin.

HeLa cells were cultured in Eagle’s minimal essential medium supplemented with 10% FBS at 37°C and 5% CO_2._ HeLa cells were treated with 100nm insulin and EGF for 30 minutes. β5 blocking (5 μg/mL) antibody or a control antibody was administered 1 h before insulin or EGF treatment.

### Primary myoblast isolation

Primary myoblast from Hind leg skeletal muscles of WT and Ptp1b KO mice were isolated following previously published protocol (5). Cells were cultured on matrigel pre-coated dish at 37ºC CO_2_ incubator and passaged when at 70% confluence. Cells were pre-plated on non-matrigel-coated dishes for 1 h after each passage to remove non-myoblast population. To differentiate myoblasts into myotubes, myoblasts were cultured to 90% confluency and subsequently cultured in differentiation media (2% horse serum and 1% Pen-Strep in DMEM) for 3-4 days.

### Protein isolation and Western blot

For protein isolation, cells were scraped in PBS, washed and centrifuged. Proteins from Cell pellets were isolated using protein isolation buffer (20mM Tris-HCl pH8.0, 137mM NaCl, 1% Nonidet P-40 (NP-40) and 2mM EDTA). To isolate protein from skeletal muscle, tissues were chopped into pieces and washed in cold PBS 2 times. Tissues were then lysed in protein isolation buffer using a tissue-lyser (Qiagen). Protein concentration was measured by Bradford assay. 20-40 μg protein samples were resolved by SDS-PAGE in 7.5% gels (Bio-Rad) and transblotted onto polyvinylidene fluoride membranes (Millipore). Membranes were blocked with 5% BSA-TBST for 1 hour and then incubated with primary antibody overnight at 4°C. Membranes were incubated with HRP-conjugated secondary antibodies for 1 hour, washed and bands were generated using an Immobilon Western chemiluminescence HRP-conjugated substrate (Amersham) and developed on a film (Kodak). Membranes were de-probed using Restore western blot stripping buffer (Thermo scientific) and reprobed for other primary antibodies.

### Co-immunoprecipitation

Proteins were isolated in non-denaturing lysis buffer (20mM Tris-HCl pH8.0, 137mM NaCl, 1% Nonidet P-40 (NP-40) and 2mM EDTA) and then precleared using fast-flow protein G Dyna beads. 200-300 μg precleared proteins were incubated overnight with the primary antibodies against PTP1B or β5 integrin or a non-specific IgG control antibody. The immunoprotein complex was then attached to protein G Dyna beads followed by protein elution from the beads in 1% SDS buffer by heating at 60°C. Equal volumes of the eluates were used for immunoblotting using antibodies against IRβ, β5 integrin and PTP1B. Proteins immunoprecipitated using non-specific IgG (NS IgG) were negative control for specific antibodies. Immunoblotting with total protein lysates was used as input samples. One of the input samples was run on the same gel with the immunoprecipitated proteins to use it as a positive control.

### PTP1B phosphatase activity assay

PTP1B phosphatase activity assay was performed using a tyrosine phosphatase assay system (Promega). Endogenous phosphates were removed from the tissue extracts or cell lysates using sephadex spin column which were subsequently incubated and immunoprecipitated with an antibody targeting PTP1B. A reaction mix was then prepared using a tyrosine phosphopeptide substrate (ENDp(Y)INASL). 2 μg of immunoprecipitated protein was incubated with the substrate for 20 minutes at 37°C in a 96-well plate. The reaction was terminated using molybdate dye/additive mixture for 15 minutes at room temperature. Optical density of the samples was measured using a plate reader at 630 nm. Phosphatase activity was calculated using the phosphatase assay standard supplied with the kit.

### Glucose uptake assay

The 2NDBG uptake assay in C2C12 myotubes and mouse primary skeletal myotubes was performed using a commercial assay kit (Cayman chemicals) following previously published protocol (5). Cells were pre-incubated with insulin (100 nM) for 10 minutes followed by the addition of nonhydrolyzable fluorescent glucose analog 2NBDG (10 mg/mL) for 30 minutes in the same media. Fluorescent intensity of cellular 2NBDG (excitation: 488 nm and emission: 535 nm) were measured using a plate reader. Fluorescent intensities of 2NBDG were then normalized to wheat germ agglutinin (WGA 680) staining intensities.

### Glucose tolerance test (GTT)

6 to 8-week-old male WT and *Ptp1b* KO mice were fasted for 5 h and then injected IP with 2 gm/kg glucose. Mice were treated with IP β5 blocking antibody (5 mg/kg) (clone ALULA, provided by A. Atakilit, University of California San Francisco [UCSF]) or mouse isotype control antibody (clone 2H6-C2, ATCC CRL-1853) for 1 h before glucose administration.

### In vivo treatment with β5 blocking antibody

Mice were fasted for 4 h before injecting them IP with β5 blocking (5 mg/Kg) or isotype control antibody for 1 h. Mice were then treated with IP insulin (1 U/kg) for 15 or 60 min before harvesting the hind-leg skeletal muscle tissues (5).

### Fasting and Refeeding of Mice

Mice were fasted for 16 h followed by refeeding with normal chow diet for 1 h prior to obtaining skeletal muscle tissue samples. Mice had free access to the food during the refeeding period.

### Human Subjects

All subjects signed consent forms to participate in the study, which was approved by the University of California San Francisco (UCSF) Institutional Review Board. Subjects were part of a multiethnic clinical cohort, termed the Inflammation, Diabetes, Ethnicity, and Obesity (IDEO), consisting of 25-to 65-year-old healthy men and women living in the San Francisco Bay Area, and recruited from medical and surgical clinics at the UCSF and the Zuckerberg San Francisco General Hospital, or through local public advertisements. The subjects covered a wide range of body mass index (BMI 18.5–52 kg/m2). Individuals were excluded for smoking, not being weight-stable for the last 3 months (change >3%), having any acute or chronic inflammatory or infectious disease, liver failure, renal dysfunction, cancer, or alcohol consumption >20g per day. IDEO collects demographic, medical, dietary, and lifestyle data from all subjects using validated questionnaires.

### Anthropometric and Body Composition Measurements

Height and weight were measured using a standard stadiometer and scale, with BMI (kg/m^2^) was calculated from two averaged measurements. Waist and hip circumferences (to the nearest 0.5 cm) were measured using a plastic tape meter at the level of the umbilicus and of the greater trochanters, respectively, and waist-to-hip ratio (WHR) was calculated. Blood pressure was measured with a standard mercury sphygmomanometer on the left arm after at least 10 minute of rest. Mean values were determined from two independent measurements. Blood samples were collected after an overnight fast and analyzed for plasma glucose, insulin, serum total cholesterol, HDL-cholesterol and triglycerides (LDL-cholesterol was estimated according to the Friedwald formula). Insulin resistance was estimated by the homeostatic model assessment of insulin resistance (HOMA-IR) index calculated from fasting glucose and insulin values (36) . Subjects taking insulin were excluded from HOMA-IR analyses. Body composition of the subjects was estimated by dual-energy X-ray absorptiometry (DXA) using a Hologic Horizon/A scanner (3-minute whole-body scan, <0.1 G mGy) per manufacturer protocol. A single technologist analyzed all DXA measurements using Hologic Apex software (13.6.0.4:3) following the International Society for Clinical Densitometry guidelines. Visceral adipose tissue (VAT) was measured in a 5-cm-wide region across the abdomen just above the iliac crest, coincident with the fourth lumbar vertebrae, to avoid interference from iliac crest bone pixels and matching the region commonly used to analyze VAT mass by CT scan (37).

### Human data statistical analysis

Normally distributed data were presented as mean +/-standard deviation for continuous measures and categorical data were expressed percentages. Unpaired independent Student’s t-test was used to compare the differences between the two groups. One way analysis of variance was used to compare differences between groups. Categorical variables were compared using the Chi-squared test. Spearman rank correlation analysis was used to determine relationships among the independent variables and HOMA-IR. Multivariate linear regression models were fit for HOMA-IR as the dependent variable and only variables significantly related (p<0.05) to HOMA-IR by Spearman correlation were entered in the linear regression analysis. A two-sided value of p<0.05 was considered statistically significant. All statistical analysis of the human data was performed using STATA version 15.1 (STATCorp LLC. College Station, TX).

### Statistical analysis

One-way ANOVA was used to compare data between multiple groups followed by Bonferroni’s posttest. For analysis of blood glucose levels over time during GTTs, a two-way ANOVA for repeated measures followed by Bonferroni’s posttest were used. All statistical analysis was performed using GraphPad Prism 9.0. Data are represented as mean ± SEM.

### Study Approval

All experiments were approved by the Institutional Animal Care and Use Committee of UCSF and the UCSF Institutional Review Board.

## Acknowledgement

This work was supported by awards from the NIH (DK110098, KA), the National Center for Advancing Translational Sciences (UCSF-CTSI UL1 TR001872, DLA), Robert Wood Johnson Foundation Harold Amos Mentored Career Award (DLA), the American Diabetes Association (1-18-PMF-003, DLA), and the San Francisco Department of Medicine Award for Excellence in Cohort Development (SK). We thank Amha Atakilit and Dean Sheppard for providing the β5 integrin blocking antibody. We thank S. Layer for ongoing inspiration.

---

**Supplementary table 1:** Baseline Characteristics by BMI class. Data are presented as mean (SD) for continuous measures, and n (%) for categorical measures.

|  | Lean (BMI≤25) (N=33) | Overweight/Obese (BMI>25) (N=56) | P-value |
| --- | --- | --- | --- |
| Male % (N) | 30.3 (10) | 48.2 (27) | 0.098 |
| Age, years | 41.67 (12.81) | 47.95 (10.99) | 0.016 |
| Ethnicity |  |  | 0.24 |
| White | 12 (36.36%) | 19 (33.93%) |  |
| Chinese | 15 (45.45%) | 18 (32.14%) |  |
| Hispanic | 6 (18.18%) | 19 (33.93%) |  |
| Weight, Kg | 61.07 (10.47) | 96.56 (24.32) | <0.001 |
| BMI | 21.97 (2.22) | 36.22 (8.18) | <0.001 |
| %Body Fat | 30.87 (6.06) | 39.37 (7.49) | <0.001 |
| Systolic BP | 123 (15.41) | 133.39 (13.11) | 0.001 |
| Diastolic BP | 71.91 (15.79) | 80.03 (10.81) | 0.006 |
| Hemoglobin A1c | 5.52 (0.66) | 6.53 (1.58) | 0.005 |
| Total Cholesterol, mg/dL | 189.55 (28.94) | 191.43 (49.31) | 0.84 |
| Triglyceride, mg/dL | 90.73 (50.37) | 148.91 (169.49) | 0.058 |
| LDL, mg/dL | 112.88 (28.08) | 115.31 (41.60) | 0.77 |
| HDL, mg/dL | 58.45 (13.28) | 49.00 (14.15) | 0.002 |
| HOMA-IR | 2.10 (1.79) | 9.27 (12.25) | 0.001 |
| Insulin, mU/L | 7.93 (3.80) | 28.47 (27.60) | <0.001 |
| Fasting glucose, mg/dL | 95.24 (16.67) | 117.41 (45.34) | 0.008 |
| Serum Mfge8, pg/mL | 2480.45 (2163.26) | 2463.58 (1809.93) | 0.97 |
| Insulin use (%) | 0 | 10 | 0.010 |
| Presence of T2D (%) | 1 (3.03) | 23 (41.07) | <0.001 |

**Supplemental Table 2:**
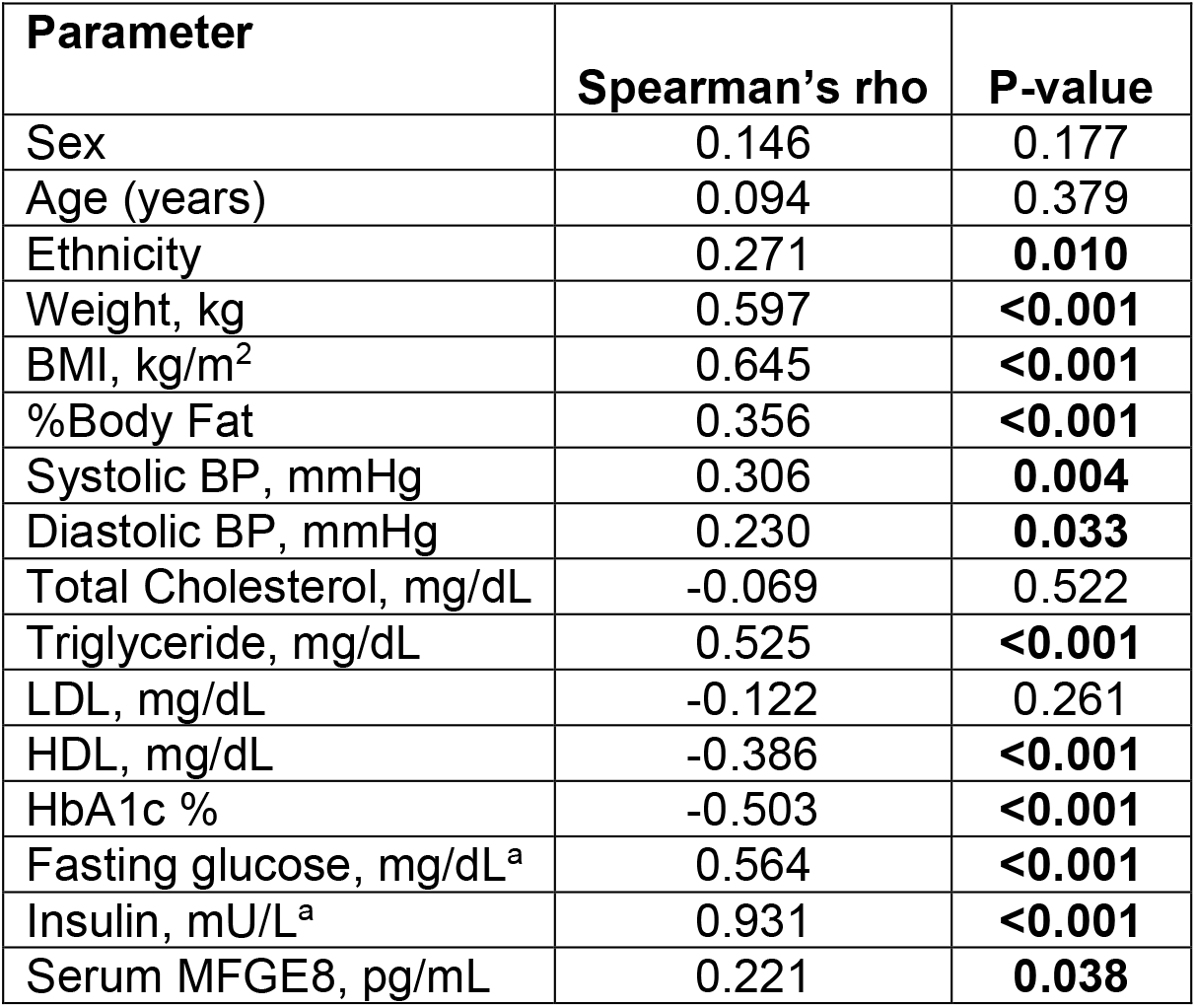
Correlation between HOMA-IR levels with demographic, body composition and biochemical characteristics. The relationships between HOMA-IR and various parameters were investigated using Spearman rank correlation. BMI, body mass index; BP, blood pressure; LDL, low-density lipoprotein cholesterol; HDL, high-density lipoprotein cholesterol; HbA1c, glycosylated hemoglobin; HOMA-IR (homeostasis model assessment of insulin resistance) = fasting insulin (mIU/L) × [fasting glucose (mg/dL)/405]; MFGE8, milk fat globule-epidermal growth factor. ^a^ Subjects on insulin were excluded from the analysis. Boldface *P* values are statistically significant (*P* < 0.05)

**Supplementary Table 3:** Determinants of HOMA-IR in multivariate regression analysis. Multivariate Linear regression analysis for the association of HOMA-IR as a dependent variable. BMI, body mass index; BP, blood pressure; HDL, high-density lipoprotein cholesterol; HbA1c, Hemoglobin A1c; HOMA-IR, homeostasis model assessment of insulin resistance; MFGE8, milk fat globule-epidermal growth factor. Boldface *P* values are statistically significant (*P* < 0.05)

| Variable | B coefficient | 95% CI | p- value |
| --- | --- | --- | --- |
| Ethnicity |  |  |  |
| Chinese | 0.9953148 | .6899138,1.435906 | 0.980 |
| Hispanic | 1.008848 | .6662883,1.527528 | 0.966 |
| Weight, kg | 1.00279 | .9871913,1.018635 | 0.723 |
| BMI, kg/m<sup>2</sup> | 1.006808 | .9598076,1.05611 | 0.777 |
| %Body Fat | 1.00497 | .9773017,1.033421 | 0.723 |
| Systolic BP, mmHg | 0.9943581 | .9822181,1.006648 | 0.359 |
| Diastolic BP, mmHg | 1.012484 | .9979501,1.02723 | 0.091 |
| HbA1c % | 1.004232 | .8610519,1.171221 | 0.956 |
| Triglyceride, mg/dL | 1.000612 | .9996838,1.00154 | 0.192 |
| HDL, mg/dL | 1.00158 | .9901064,1.013187 | 0.784 |
| Fasting blood glucose, mg/dL | 1.009417 | 1.003248,1.015625 | **0.003** |
| Fasting Insulin, mU/L | 1.024313 | 1.015021,1.03369 | **<0.0001** |
| Serum MFGE8, pg/mL | 1.000079 | 1.000004,1.000153 | **0.040** |

